# MEXCOwalk: Mutual Exclusion and Coverage Based Random Walk to Identify Cancer Modules

**DOI:** 10.1101/547653

**Authors:** Rafsan Ahmed, Ilyes Baali, Cesim Erten, Evis Hoxha, Hilal Kazan

## Abstract

**Motivation:** Genomic analyses from large cancer cohorts have revealed the mutational heterogeneity problem which hinders the identification of driver genes based only on mutation profiles. One way to tackle this problem is to incorporate the fact that genes act together in functional modules. The connectivity knowledge present in existing protein-protein interaction networks together with mutation frequencies of genes and the mutual exclusivity of cancer mutations can be utilized to increase the accuracy of identifying cancer driver modules.

**Results:** We present a novel edge-weighted random walk-based approach that incorporates connectivity information in the form of protein-protein interactions, mutual exclusivity, and coverage to identify cancer driver modules. MEXCOwalk outperforms several state-of-the-art computational methods on TCGA pancancer data in terms of recovering known cancer genes, providing modules that are capable of classifying normal and tumor samples, and that are enriched for mutations in specific cancer types. Furthermore, the risk scores determined with output modules can stratify patients into low-risk and high-risk groups in multiple cancer types. MEXCOwalk identifies modules containing both well-known cancer genes and putative cancer genes that are rarely mutated in the pan-cancer data. The data, the source code, and useful scripts are available at: https://github.com/abu-compbio/MEXCOwalk.

## 1 Introduction

Recent advances in high-throughput DNA sequencing technology have allowed several projects such as the TCGA (Weinstein *et al*., 2013) to construct and release genomic data from thousands of tumors. This further gave rise to the design of several computational approaches for the systematic detection of cancer-related somatic mutations.

Combining the somatic mutations data with additional information in the form of interaction networks or gene expression data, several computational approaches focus on prioritizing independent genes to provide hypothesized *candidate driver genes*, those defined as being causally linked to oncogenesis (Erten *et al*., 2011; Lawrence *et al*., 2013; Yang *et al*., 2017b; Dopazo and Erten, 2017). Although such gene rankings provide valuable insight regarding potential genes of interest, in many cases mutations at different loci could lead to the same disease (Vanunu *et al*., 2010). This genetic heterogeneity may reflect an underlying molecular mechanism in which the cancer-causing genes form some kind of functional pathways or *candidate driver modules*. Several computational methods have been suggested for the identification of candidate modules; see Dimitrakopoulos and Beerenwinkel (2017); Deng *et al*. (2019); Zhang and Zhang (2018) for recent surveys.

The module identification approaches as applied to cancer can be viewed in two broad categories based on the types of input data they employ. The *de novo* methods rely only on genetic data to discover novel genetic interactions, as well as cancer-related functional modules (Miller *et al*., 2011; Vandin *et al*., 2011b; Leiserson *et al*., 2013; Liu *et al*., 2017). Due to the large solution space such methods usually apply a prefiltering based on alteration frequency to reduce the inherent computational complexity which may reduce sensitivity by overlooking modules involving rare alterations (Deng *et al*., 2019).

On the other hand, knowledge-based methods, in addition to genomic data, incorporate prior knowledge in the form of pathways, networks and functional phenotypes to identify driver modules. Such methods can be subcategorized based on the optimization goals set within the computational problem formulations they employ in defining the biologically motivated cancer driver module identification problem. Since a driver pathway tends to be perturbed in a relatively large number of patients, in the first subcategory of methods including Hotnet (Vandin *et al*., 2011a), Hotnet2 (Leiserson *et al*., 2014), Hierarchical Hotnet (Reyna *et al*., 2018), the *coverage* of the modules as identified by the mutation frequencies of the comprising genes over a cohort of samples constitutes an informal optimization goal. Heat-diffusion over an interaction network that diffuses the mutation frequencies throughout the network is a common attribute in these methods. The resulting diffusion values are then employed to extract modules exhibiting a large degree of connectedness as formulated with an appropriate graph-theoretical connectivity definition, usually the *strong connectivity*. The second subcategory of knowledge-based module identification methods incorporate an appropriate definition of an important concept, *mutual exclusivity*, in their computational problem formulations. It refers to the phenomenon that for a group of genes which exhibit evidence of shared functional pathway, simultaneous mutations in the same patients are less frequent than is expected by chance (Yeang *et al*., 2008). Several cancer module identification methods incorporate this observation in the employed combinatorial optimization problem definitions. In MEMo, maximal clique extraction in a similarity graph derived from an interaction network or functional relation graph is used and the maximal cliques are postprocessed taking into account the mutual exclusivity results (Ciriello *et al*., 2012). In Babur *et al*. (2015) a method based on seed-and-growth on a network, where the growth strategy is determined with respect to a suitably defined mutual exclusion score is proposed to identify pan-cancer modules using TCGA data. BeWith proposes an ILP formulation that combines interaction density within a module and several mutual exclusivity definitions as a maximization goal (Dao *et al*., 2017). The ILP formulation incorporates constraints in the form of desired number of modules and maximum number of genes per module. MEMCover combines pairwise mutual exclusion scores with confidence values of interactions in the network (Kim *et al*., 2015). To maximize high-confidence interactions, mutual exclusivity, and coverage simultaneously; heavy subnetworks covering every disease case at least *k* times are found following a greedy iterative seed-and-growth heuristic.

We propose MEXCOwalk, a knowledge-based method that incorporates protein-protein interaction (PPI) network data and mutation profiles, and employs a random walk-based approach to extract driver modules for cancer. We first provide a novel optimization problem definition for identifying driver modules, that takes into account network connectivity, mutual exclusivity, and coverage. Computational intractability of the provided optimization problem is shown for completeness. MEXCOwalk is inspired by the Hotnet2 method and its variants, and extends them in two important aspects. Firstly, similar to Hotnet2 we create a vertex-weighted graph to apply random-walk on, where vertex weights correspond to coverages. However, different from Hotnet2, our graph is also edge-weighted, where the edge weights reflect a novel combination of the coverages and the degree of mutual exclusivity between pairs of gene neighborhoods. To our knowledge, this is the first method to employ edge-weighted random walks for identifying driver modules. Secondly, we provide a novel heuristic based on split-and-extend, where certain modules are split into pieces to be recombined into new modules while maintaining high coverage and mutual exclusivity. We show that MEXCOwalk provides better results than three alternative knowledge-based methods in terms of recovering known cancer genes including the rarely mutated ones, enrichment for mutations in specific cancer types, and the accuracy in classifying normal and tumor instances.

## 2 Methods

In the following subsections we provide the problem definition and a description of our MEXCOwalk algorithm.

### 2.1 Problem Definition

We provide a novel combinatorial optimization problem definition to detect driver modules in cancer. Such a definition is not only important for algorithmic purposes but also to serve as a measure of performance for alternative methods suggested for the problem.

Let *S_i_* denote the set of samples for which gene *g_i_* is mutated. Let *G* = (*V, E*) represent the PPI network where each vertex *u_i_* ∈ *V* denotes a gene *g_i_* whose expression gives rise to the corresponding protein in the network and each undirected edge (*u_i_, u_j_*) ∈ *E* denotes the interaction among the proteins corresponding to genes *g_i_, g_j_*. Henceforth we assume that *g_i_* denotes both the gene and the corresponding vertex in *G*.

Let *M* ⊆ *V* be a set of genes denoting a *module*. We define the mutual exclusivity of *M* as, 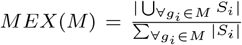 and the coverage of *M* as, 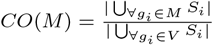. We note that although such definitions have been employed in previous work, the module sizes have not been taken into consideration (Wu *et al*., 2015, 2016).

Let *P* = (*M*_1_, *M*_2_,… *M_r_*} be a set of modules. Let *RS*(*M_q_*) denote the relative size of a module *M_q_* with respect to the total size, that is 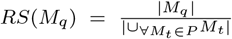. We define the mutual exclusivity score and the coverage score of a set of modules, so that each module *M_q_* contributes its share proportional to its relative size *RS*(*M_q_*) for the former, whereas for the latter the contribution of Mq is proportional to the normalized value of 1 − *RS*(*M_q_*). Intuitively, a large module with high mutual exclusivity score should be rewarded, since as the size of the module increases the chances of achieving better mutual exclusivity decrease. Analogously, a small module with high coverage score should be rewarded. Thus we define the mutual exclusivity score of *P* as, 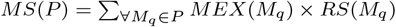. The coverage score of *P* is defined as 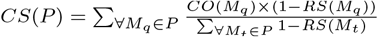 if |*P*| > 1 and *CS*(*P*) = *CO*(*M*_1_), if |*P*| = 1.

For a graph *H* and a set *M_q_* of genes, let *H*(*M_q_*) denote the subgraph of *H* induced by the vertices corresponding to genes in *M_q_*.

#### Cancer driver module identification problem

Given as input a PPI network *G, S_i_* for each gene *g_i_*, integers *total_genes*, and *min_module_size*, find a disjoint set of modules *P* that maximizes the *driver module set score* defined as,

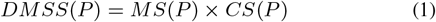

and that satisfies the following:

1. For each *M_q_* ∈ *P, G*(*M_q_*) is connected.
2. |∪_∀*M_q_*∈*P*_*M_q_*| = *total_genes*.
3. *min_∀M_q_∈P_*|*M_q_*| = *min_module_size*.

##### Theorem 1.

*Cancer driver module identification problem is NP-hard*.

Proof. See Supplementary Document.

### 2.2 MEXCOwalk Algorithm

Due to the computational intractability of the problem, we propose a polynomial-time heuristic approach. The pseudocode is provided in Algorithm 1. There are three main steps of the algorithm, each of which is described in detail in the following subsections.

#### 2.2.1 Weight Assignment with MEX and CO

Given a PPI network *G* = (*V, E*), we first construct a weighted graph *G_w_* that contains properly defined weights for vertices and edges. For each *g_i_* ∈ *V* we assign a weight, *w*(*g_i_*) = *CO*({*g_i_*}), thus the weight corresponds to the mutation frequency of a gene. It represents the heat to be diffused from that vertex during the random walk procedure.

The weight of an edge incident on a *g_i_* should on the other hand reflect the ratio of heat transferred to *g_i_*’s neighbors at each step of the random-walk. We formulate it so as to mimic the optimization goal defined in the problem definition. One option could be to define the weight solely in terms of the gene pair *g_i_, g_j_*. However such a simple weighting scheme may not be sufficient in practice, since the co-occurrence of a pair in a module increases the chances of the genes in their neighborhoods to coexist in the same module as well. This is especially important for the contribution of mutual exclusivity in the edge-weight, as pairwise mutual exclusivity values are almost always close to 1. In order to reflect these observations we consider an edge-weighting scheme where contribution of mutual exclusivity is computed within the vertex neighborhoods. More specifically, let *N_e_*(*g_i_*) denote the *extended neighborhood* of *g_i_*, that is *N_e_*(*g_i_*) = ∪_∀(*g_i_, g_j_*)∈*E*_ *g_j_* ∪ {*g_i_*}. The contribution of mutual exclusivity to the edge weight, *MEX_n_*(*g_i_, g_j_*) is the average of MEX(*N_e_*(*g_i_*)) and *MEX*(*N_e_*(*g_j_*)). Thus the weight of an edge (*g_i_, g_j_*) is defined as, *w*(*g_i_, g_j_*) = *MEX_n_*(*g_i_, g_j_*) × *CO*({*g_i_*}) × *CO*({*g_j_*}). The contribution of coverage is computed as a product so as to reduce the chances of a single gene with large coverage dominating the weights of incident edges. Furthermore, it allows the algorithm to favor more balanced coverages among equal-sized modules; coverage of 100 patients with a module containing a pair of genes, one covering 99 and the other only 1, is less preferable than a module with a pair where each gene covers 50 patients. Finally we note that, to further strengthen the impact of mutual exclusivity on edge-weights, we introduce athreshold *θ*, so that for pairs with *MEX_n_* score less than *θ*, edge weights are assigned to 0.

#### 2.2.2 Edge-Weighted Random Walk

Once *G_w_* is constructed after vertex and edge weight assignments, we apply an insulated heat diffusion process on *G_w_* that can also be described as a random walk with restart on the graph. The random walk starts from a gene *g*_s_. At each time step, with probability 1 − *β*, the random surfer follows one of the edges incident on the current node with probability proportional to the edge weights. Otherwise, with probability *β*, the walker restarts the walk from *g_s_*. Here *β* is called the restart probability. The transition matrix *T* corresponding to this process can be constructed by setting 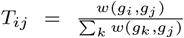, if (*g_i_, g_j_*) ∈ *E*, and *T_ij_* = 0 otherwise. Thus *T_ij_* can be interpreted as the probability that a simple random walk will transition from *g_j_* to *g_i_*. The random walk process can then be described as a network propagation process by the equation, *F*_*t*+1_ = (1 − *β*)*TF_t_* + *βF*_0_, where *F_t_* is the distribution of walkers after *t* steps and *F*_0_ is the diagonal matrix with initial heat values, that is *F*_0_ [*i, i*] = *CO*(*g_i_*). One strategy to compute the final distribution of the walk is to run the propagation function iteratively for increasing *t* values until *F*_*t*+1_ converges (Hofree *et al*., 2013). Another strategy, which we chose to employ in our implementation, is to solve this system numerically using the equation, *F* = *β*(*I* − (1 − *β*)*T*)^−1^*F*_0_ (Leiserson *et al*., 2014). The edge-weighted directed graph *G_d_* is constructed by creating directed edge [*g_i_, g_j_*] with weight *F*[*i, j*], for every pair *i* ≠ *j*.

The idea of random walks with restart has been employed in the context of cancer module identification in previous work (Vandin *et al*., 2011a; Leiserson *et al*., 2014; Bersanelli *et al*., 2016; Yang *et al*., 2017a; Reyna *et al*., 2018). However as the concept of edge weights is absent, the transition probabilities in those studies are only based on the degrees of the vertices. In our case, the transition probabilities reflect the edge weights which in turn model the contribution of a pair of genes to the maximization score, when placed in the same module. Similar to the previous methods employing heat diffusion we assign *β* = 0.4.

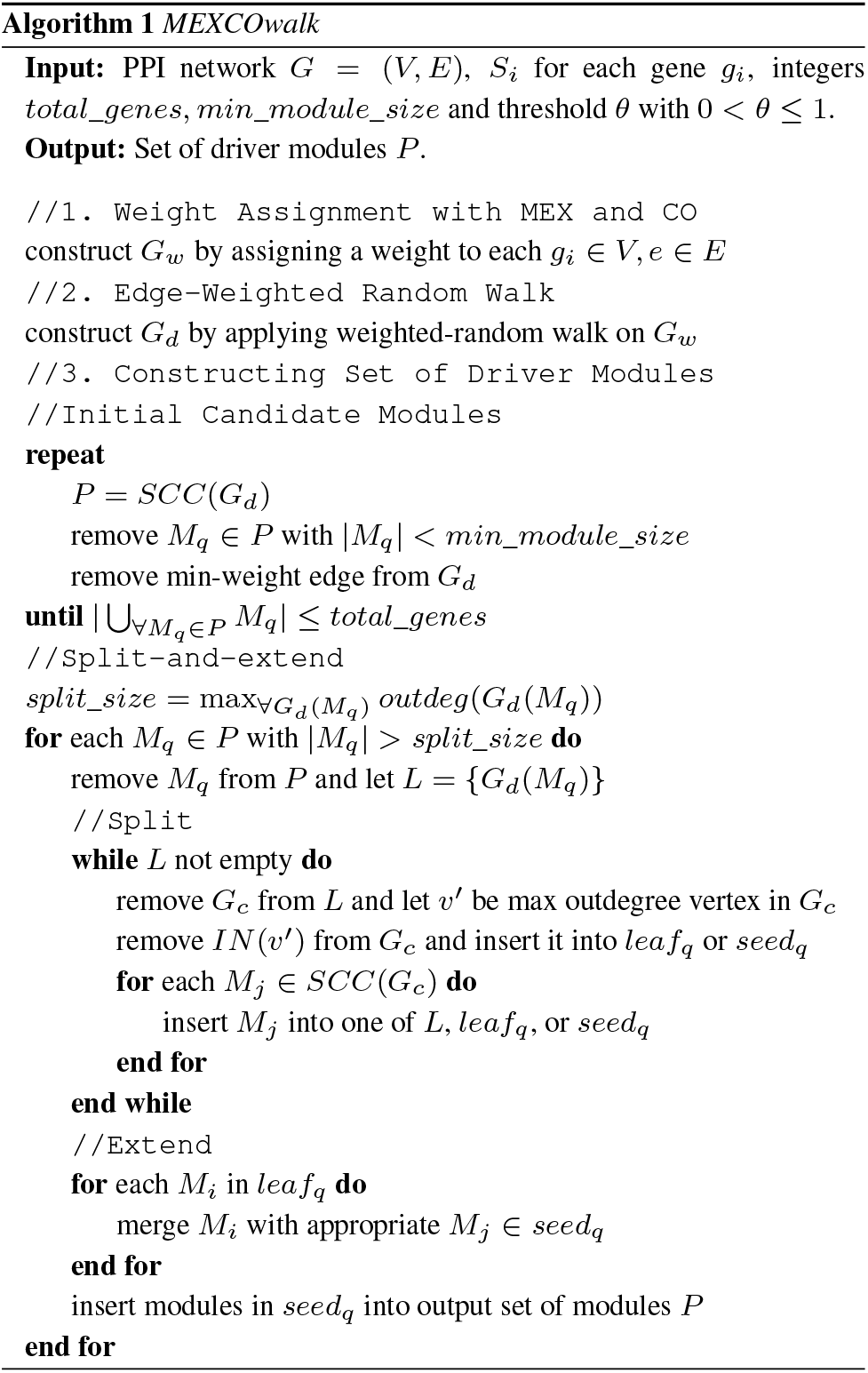

#### 2.2.3 Constructing Set of Driver Modules

We have two main steps. We employ strongly connected components (SCC) as a primitive in both of the steps. We first create an initial set of candidate modules. For this, we iteratively remove the smallest weight edge from *G_d_*, add the strongly connected components (SCC) of *G_d_* into initial module set *P*, and remove all modules of size less than *min_module_size* from *P*, until the total number of genes in *P* decreases to *total_genes*. The idea of employing SCCs is inspired by Hotnet2. However, for Hotnet2 the SCCs comprise the final set of modules, whereas we further process the SCCs via a novel *split-and-extend* procedure. The aim of this procedure is to split modules larger than a certain size into pieces that can be recombined with respect to degrees of connectivity in *G_d_*, which in turn correspond to the achieved mutual exclusivity and coverage via the edge weights. We define the *split_size* to be the outdegree of any vertex in any of the subgraphs induced by the modules. Any initial candidate module *M_q_* of size greater than the *split_size* goes through the split-and-extend procedure. The idea is to first extract *seed* modules that satisfy certain size and connectivity criteria, and extend them with small *leaf* modules. Given a directed graph *G_c_*, let *IN*(*v*′) denote the *isolated neighborhood* of *v*′ in *G_c_*, that is *w* ∈ *IN*(*v*′), if and only if *w* ∈ *N_e_*(*v*′) and for any directed edge [*w, x*] or [*x, w*], *x* ∈ *N_e_*(*v*′).The split phase of a module *M_q_* consists of removing *IN*(*v*′) from *G_d_*(*M_q_*), where *v*′ is the vertex with largest degree in *G_d_*(*M_q_*). Assuming its size is not less than *min_module_size, IN*(*v*′) is a seed module to be extended in the next phase, otherwise it is a leaf module that is to be attached to an appropriate seed module. The remainder of *G_d_*(*M_q_*) goes through a SCC partitioning. Any resulting component of size larger than the split_size goes through the same split process, any component of size less than *min_module_size* becomes a leaf module, and any other component in between these two sizes becomes a seed module. In the extend phase, each leaf module is merged with the seed module with which it has maximum number of connections in *G_d_*(*M_q_*).

## 3 Discussion of Results

We implemented the MEXCOwalk algorithm in Python. The source code, useful scripts for evaluations, and all the input data are freely available as part of the supplementary material. We compare MEXCOwalk results against those of three other existing knowledge-based cancer driver module identification methods: Hotnet2, MEMCover, and Hierarchical Hotnet. The first two benchmark algorithms are chosen as representatives of their respective subcategories; Hotnet2 is a popular benchmark method based on optimizing coverage via a heat-diffusion heuristic and MEMCover is a popular algorithm among those optimizing mutual exclusivity as well as coverage via a greedy seed-and-growth heuristic. Hierarchical Hotnet is chosen as a third benchmark method, as it is one of the most recent cancer driver module identification methods.

### 3.1 Input Data and Parameter Settings

All four methods, including MEXCOwalk, assume same type of input data in the form of mutation data of available samples and a H.Sapiens PPI network. We employ somatic aberration data from TCGA, preprocessed and provided by Leiserson *et al*. (2014). The preprocessing step includes the removal of hypermutated samples and genes with low expression in all tumor types. After the filtering, the dataset contains somatic aberrations for 11,565 genes in 3,110 samples. The mutation frequency of a gene *g_i_* is calculated as the number of samples with at least one single nucleotide variation (SNV) or copy number alteration (CNA) in *g_i_* divided by the number of all samples. As for the PPI network, we used the HINT+HI2012 network (Yu *et al*., 2011; Das and Yu, 2012; Leiserson *et al*., 2014). We execute each of the four algorithms on the largest connected component of this combined network that consists of 40,704 interactions among 9,858 proteins.

Regarding MEXCOwalk, we have settings for three parameters: the mutual exclusivity threshold *θ*, the *total_genes*, and the *min_module_size*. In the main document, we present results for *θ* = 0.7. The results with other threshold values are available in the Supplementary Document. The *total_genes* parameter is considered the main independent variable; we obtain the results of each evaluation under the settings *total_genes =* 100, 200,…, 2500. Finally, we set *min_module_size* to 3 for the results discussed in the main document, as this constitutes a nontrivial mimimum module size compatible with the problem definition. Further results of the settings of *min_module_size* are in the Supplementary Document. For Hotnet2, we obtain results for varying values of *total_genes* = 100, 200,…, 2500, with the default value of *min_module_size* = 3. We present results of Hierarchical Hotnet where the *clustering parameter δ* is determined by the recommended permutation test. Hierarchical Hotnet outputs a total of 806 genes in modules of size greater than one. Since some of these modules may contain modules with two genes, we generate a filtered version as well, where all such modules are removed, resulting in modules with a total of 554 genes. In what follows, we refer to the former version as *HierHotnet_v1* and the latter version as *HierHotnet_v2*. For MEMCover, as recommended in the original paper, mutual exclusivity scores are obtained from type-restricted permutation test with all pancancer samples, that is the TR_test. Because confidence scores are not available for HINT + HI2012 network, we set the confidence score of all edges to 1 when calculating the edge weights for the MEMCover model. We set the coverage parameter *k* to its default value of 15. MEMCover introduces a parameter, *f*(*θ*), that is used to control the tradeoff between the output number of modules and the average weights within each module. It indirectly controls the module sizes; the smaller *f*(*θ*), the larger the modules output by MEMCover in general. We consider three settings for the MEMCover algorithm, referred to as *MEMCover_v1*, *MEMCover_v2*, and *MEMCover_v3*, respectively. For the first one, we assign *f*(*θ*) = 0.584, which is achieved by setting *θ* parameter (not to be confused with the *θ* we employ in MEXCOwalk) to 40%, as recommended in the original paper. For the second one, we assign *f*(*θ*) = 0.03, which is the setting that minimizes the percentage of size one and size two modules. Finally, the last one corresponds to the setting where *f*(*θ*) = 0.03 and all modules of size < 3 are removed. To obtain results with varying *total_genes* from 100 to 2500 we consider the modules formed by the first *total_genes* many genes output by each version, since the order MEMCover outputs the modules reflects the algorithm’s quality preferences. Values of *total_genes* larger than 1600 are not available for *MEMCover_v3* as it outputs 1684 genes in total.

### 3.2 Static Evaluations

Most of the existing driver module identification methods employ *static evaluations*, where the union of the genes in all the modules are compared against a reference set of cancer genes. For consistency with previous work, our first evaluation compares the algorithms based on their ability to recover these known cancer genes. COSMIC Cancer Gene Census (CGC) database (Forbes *et al*., 2017) is one popular reference gene set containing 616 genes with mutations that have been causally implicated in cancer. Out of 616 genes 504 are in the dataset we employ. The Area Under the ROC (AUROC) analysis with respect to the COSMIC gene set indicates that MEXCOwalk and *MEMCover_v1* have the same AUROC value of 0.083. *MEMCover_v2* ranks the second with 0.078. The AUROC value of Hotnet2 is 0.067. AUROC is undefined for *HierHotnet_v1*, *HierHotnet_v2*, and *MEMCover_v3*. Nevertheless inspecting *MEMCover_v3’s* Receiver Operating Characteristic (ROC) curve plots, we can observe that its outputs provide worse True Positive (TP) rates than those of *MEMCover_v2* and better rates than those of Hotnet2. The results of *HierHotnet* versions almost overlap with those of Hotnet2. Another reference gene set is DGIdb 3.0, which contains a set of 1062 druggable genes identified by mining existing resources on how mutated genes might be targeted therapeutically or prioritized for drug development (Coffman *et al*., 2017). With respect to this reference set, MEXCOwalk achieves the best AUROC value of 0.043, followed by *MEMCover_v1* and *MEMCover_v2*, each with an AUROC of 0.040. Finally, Hotnet2 achieves an AUROC of 0.039.

To find out the performance of the module finding algorithms in identifying genes with rare mutations, we repeat the above analysis, limiting each reference to the set of genes that have upto 1% and upto 2% mutation frequencies in the pan-cancer patient cohort under study. With regard to the COSMIC gene set, out of 504 genes, 342 are in the 1% frequency range and 438 are in the 2% frequency range. MEXCOwalk performs the best, achieving AUROC values of 0.082 and 0.085, for the frequencies of 1% and 2%, respectively. AUROC values of *MEMCover_v1, MEMCover_v2*, and Hotnet2 are respectively 0.077, 0.071, 0.069 for the 1% frequency case and 0.081, 0.074, 0.070 for the 2% frequency case. With respect to the DGIdb 3.0 reference set, out of 1062 genes, 913 are in the 1% range and 1015 are in the 2% range. MEXCOwalk again achieves the highest AUROC values of 0.044 and 0.045, for the frequencies of 1% and 2%, respectively. *MEMCover_v1* and *MEMCover_v2* both have an AUROC value of 0.041 and Hotnet2 has an AUROC value of 0.039 for both frequencies. Detailed figures plotting the ROC curves of the set of genes in the union of modules of each algorithm with respect to the CGC, DGIdb 3.0 and their rare mutation-filtered versions can be found in the Supplementary Document.

Finally, to emphasize the disease aspect of the problem that separates it from simple module identification in a given PPI network and to verify the effects of employed mutation frequencies we conduct further tests on randomized data. For this, we first assign the actual mutation frequencies to the set of mutated genes randomly. Next for each patient, we select as many genes as are mutated in the original patient data to be mutated, where the selection probability of each gene is proportional to newly assigned mutation frequencies. We execute MEXCOwalk on the generated data and repeat the static evaluations with respect to the CGC, DGIdb 3.0, and their rare mutation-filtered versions. Detailed results plotting overlaps with each reference set can be found in the Supplementary Document. As expected, the overlap ratios of the modules obtained with original data are much higher than those obtained with random mutations data.

### 3.3 Modular Evaluations

The static evaluations of the previous subsection measure the capability of an algorithm in dissecting cancer-related genes in the union of the modules it provides, without regard for the generated specific modules and their interrelations. With respect to this evaluation, for instance, for a fixed set of genes, an output placing every single gene of the set into its own module in one extreme, an output consisting of a single large module with all the genes in the set in another extreme, and every other output in between these extremes would all provide same scores. Neither extreme is suitable for the purposes of module identification. The original MEMCover, that is *MEMCover_v1*, provides outputs similar to the former extreme, where almost 70% of all output genes are in modules of size one. It produces modules of average size 1.2, for almost all values of *total_genes*, whereas average size of MEXCOwalk modules is between 6.5 and 9. This observation regarding module sizes indicates that, although the AUROC value of MEMCover_v1 with respect to the COSMIC reference set is as good as that of MEXCOwalk, the former only achieves this at the expense of providing trivial outputs with one gene or two genes in a module. Such outputs are against the very notion that each driver module should identify a functional pathway important for cancer. On the other hand Hotnet2 produces modules similar to the latter extreme; more than 60% of output genes are in a single large module between 500 and 2000 *total_genes* and this percentage gets to more than 80% for *total_genes >* 2000. Plots depicting the percentages of genes in modules of largest size, smallest size, and the average module sizes with respect to increasing *total_genes* for all algorithms under consideration can be found in the Supplementary Document. To compensate for such a drawback of static evaluations, we provide three modularity-based metrics and evaluate the output module sets of alternative methods based on these metrics.

#### 3.3.1 Driver Module Set Score

Our first modular evaluation metric is the main optimization goal of the cancer driver module identification problem, that is the driver module set scores (*DMSS*) defined in Equation 1. Fig. 1-A shows that MEXCoWalk modules have better *DMSS* values than the module sets of all the other methods. The difference is much more dramatic for smaller *total_genes* values such as 100 and 200. Those of Hierarchical Hotnet and Hotnet2 are among the worst, especially for settings of *total_genes >* 500. *MEMCover_v1* performs worse than the two other MEMCover versions, as it provides many size 1 and size 2 modules. This finding demonstrates another merit of the *DMSS* definition; if there are many small modules, assuming the mutual exclusivity does not decrease substantially by enlarging the modules, then our optimization score function prefers outputs with larger modules. Consider for instance, the following special case where we have 10 genes under consideration, each covering *x* out of a total of *y* samples. The output consisting of a set of modules each containing one gene has a *DMSS* of *x*/*y*. On the other hand, assuming a *MEX* score of *m* for every pair of genes, the output with any pair of genes per module has a *DMSS* of 2*m*^2^*x*/*y*. This implies that the latter is a more preferable module set than the former, as long as 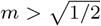. It corresponds to the case where upto almost 58% of samples covered by a gene to be in the intersection of samples covered by another gene.

**Fig. 1.**
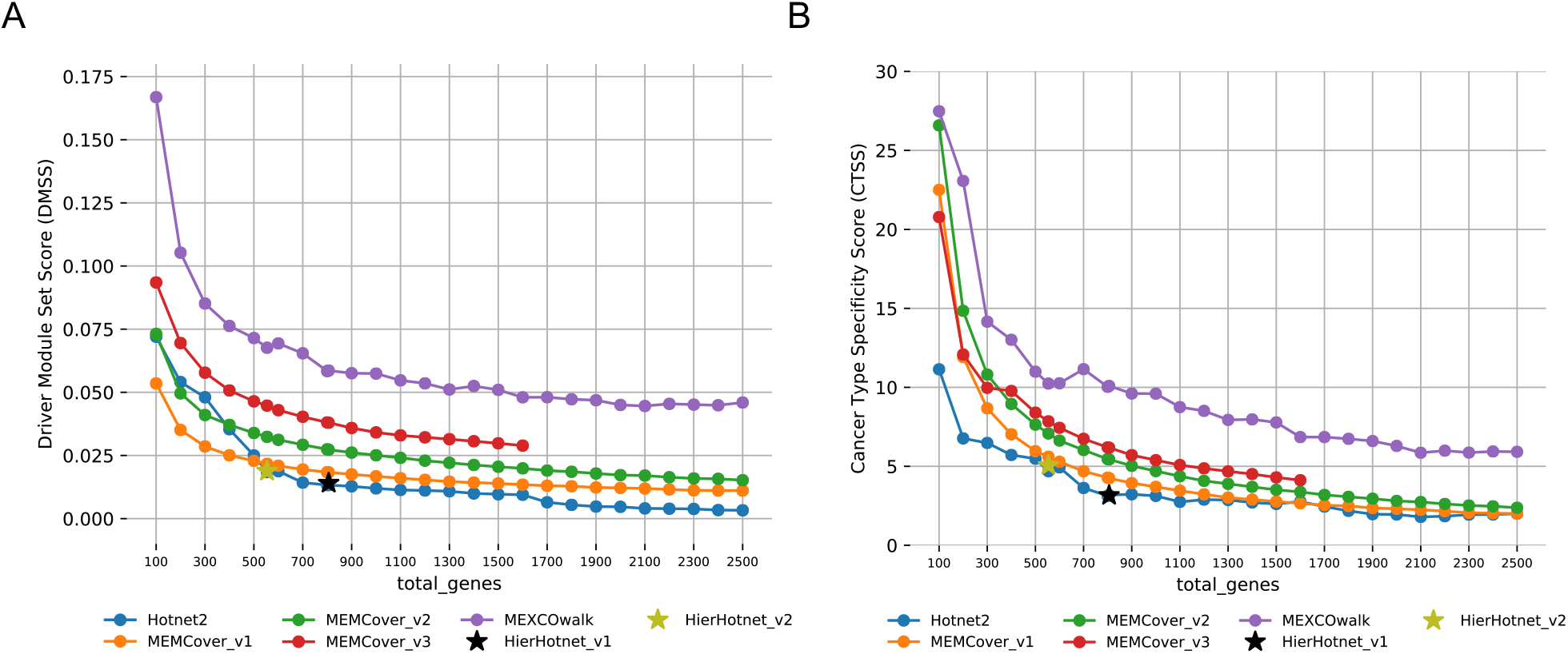
A) DMSS evaluations of output modules of MEXCOwalk, MEMCover, Hotnet2, and Hierarchical Hotnet for increasing values of *total_genes*. B) CTSS evaluations of output modules of MEXCOwalk, MEMCover, Hotnet2, and Hierarchical Hotnet for increasing values of *total_genes*.

#### 3.3.2 Cancer Type Specificity Score

Our second modularity-based evaluation metric is defined with respect to cancer type specificity. We test an output module set in terms of enrichment for mutations in a specific cancer type using Fisher’s exact test. For a module *M*, let *S_M_* denote the set of patients where at least one of the genes in *M* is mutated. For a cancer type *t*, let *S_M_^t^* denote the subset of patients in *S_M_* diagnosed with cancer type *t*. Assuming *n_t_* denotes the number of patients of cancer type *t* in the whole dataset, we calculate the Fisher’s exact test with the following entries in the contingency table in row-major order: |*S_M_^t^*|, *n_t_* − |*S_M_^t^*|, 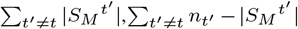. We use the False Discovery Rate (FDR) correction procedure for multiple testing correction (Benjamini and Hochberg, 1995).

Let *P* = {*M*_1_, *M*_2_,… *M_r_*} be a set of modules. For each module *M_q_* ∈ *P*, the described process results in a p-value for every cancer type *t*, denoted with *p_q_^t^*. We define the *cancer type specificity score* of *P* as the average – log of best p-value per module. More formally,

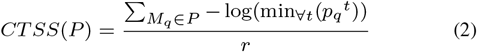

Fig. 1-B shows the *CTSS* scores of the module sets provided by the methods under consideration; see Supplementary Document for detailed distribution of individual p-values. Compared to the other methods, MEXCOwalk provides a larger *CTSS* value for every setting of *total_genes*, indicating that the output modules are strongly enriched for particular cancer types. We also observe that module sets of MEMCover versions perform better than those of Hotnet2 and Hierarchical Hotnet.

Note that Fig. 1-B, bears a striking similarity to the figure plotting *DMSS*, Fig. 1-A. This indicates that our optimization goal, as defined by the combinatorial metric *DMSS* to measure the quality of output set of modules, is further validated by a biological metric.

#### 3.3.3 Mean Classification Accuracy Score

We examine the predictive value of an output set of modules in classifying tumor and normal samples of TCGA pan-cancer data consisting of 12 cancer types. We downloaded the gene expression data of 437 normal and 4307 tumor samples from Firebrowse database (*http://firebrowse.org*; version 2016_01_28). For this, we employ k-nearest-neighbor classifier using Euclidean distance with *k* = 1, where the features are the expression values of the set of genes in a module. To evaluate the predictive performance of a module *M_q_*, we use 10-fold stratified cross-validation accuracy, denoting it with *Acc*(*M_q_, c*) for a fold *c*. We can then define the Mean Classification Accuracy Score of a set of modules *P* as,

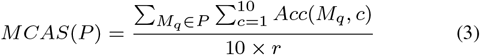

The plots of the MCAS scores of the module sets of all four methods for varying *total_genes* are provided in Fig. 2; see Supplementary Document for detailed distribution of individual accuracy values. MEXCOwalk consistently achieves the top accuracy for all settings of *total_genes*, implying that MEXCOwalk modules can more accurately perform tumor/normal classification than the other methods. Interestingly, Hierarchical Hotnet performs worse than Hotnet2. Among MEMCover models, MEMCover_v3 shows a better performance than MEMCover_v1 and MEMCover_v2, in contrast to their relative performances in recovering known cancer genes.

**Fig. 2.**
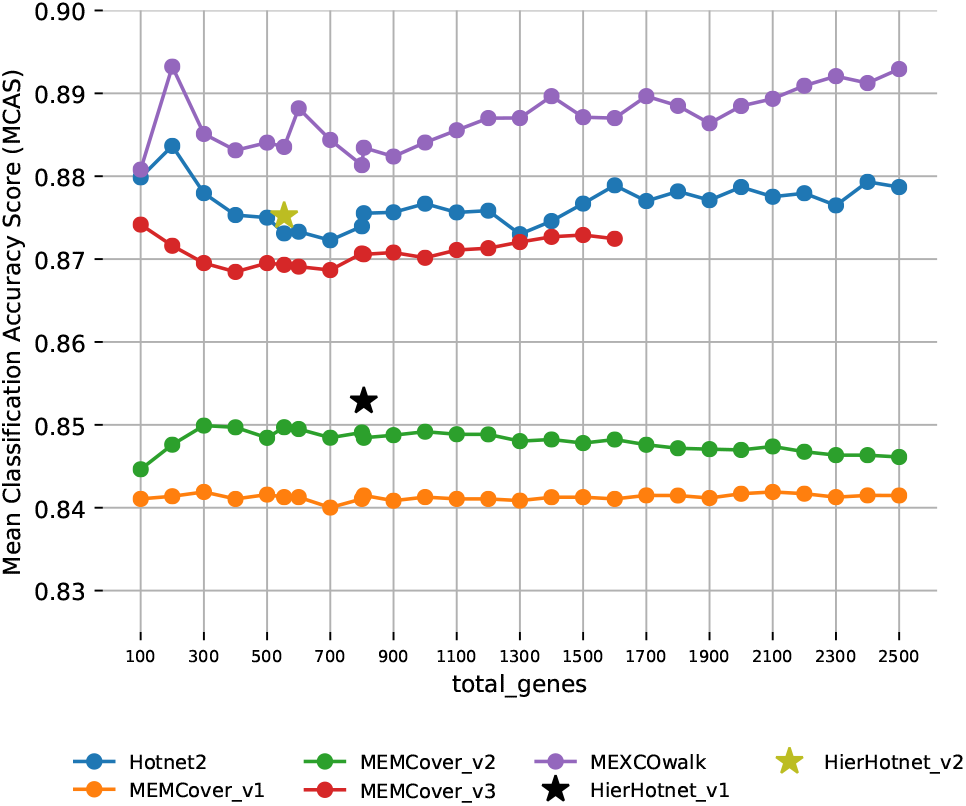
MCAS evaluations of output modules of MEXCOwalk, MEMCover, Hotnet2, and Hierarchical Hotnet for increasing values of *total_genes*.

### 3.4 Analysis of MEXCOwalk Modules

Fig. 3-A shows the 12 modules that MEXCOwalk identifies when *total_genes* is set to 100. The sizes of the modules range between 3 and 31, and their coverage values range between 5% to 50%. Note that the edges correspond to the PPI network edges, whereas the weight of an edge is the smaller of the weights of the corresponding directed edges from *G_d_*, as computed through edge-weighted random walk and thus represents the degree of mutual exclusivity and coverage assigned by MEXCOwalk.

**Fig. 3.**
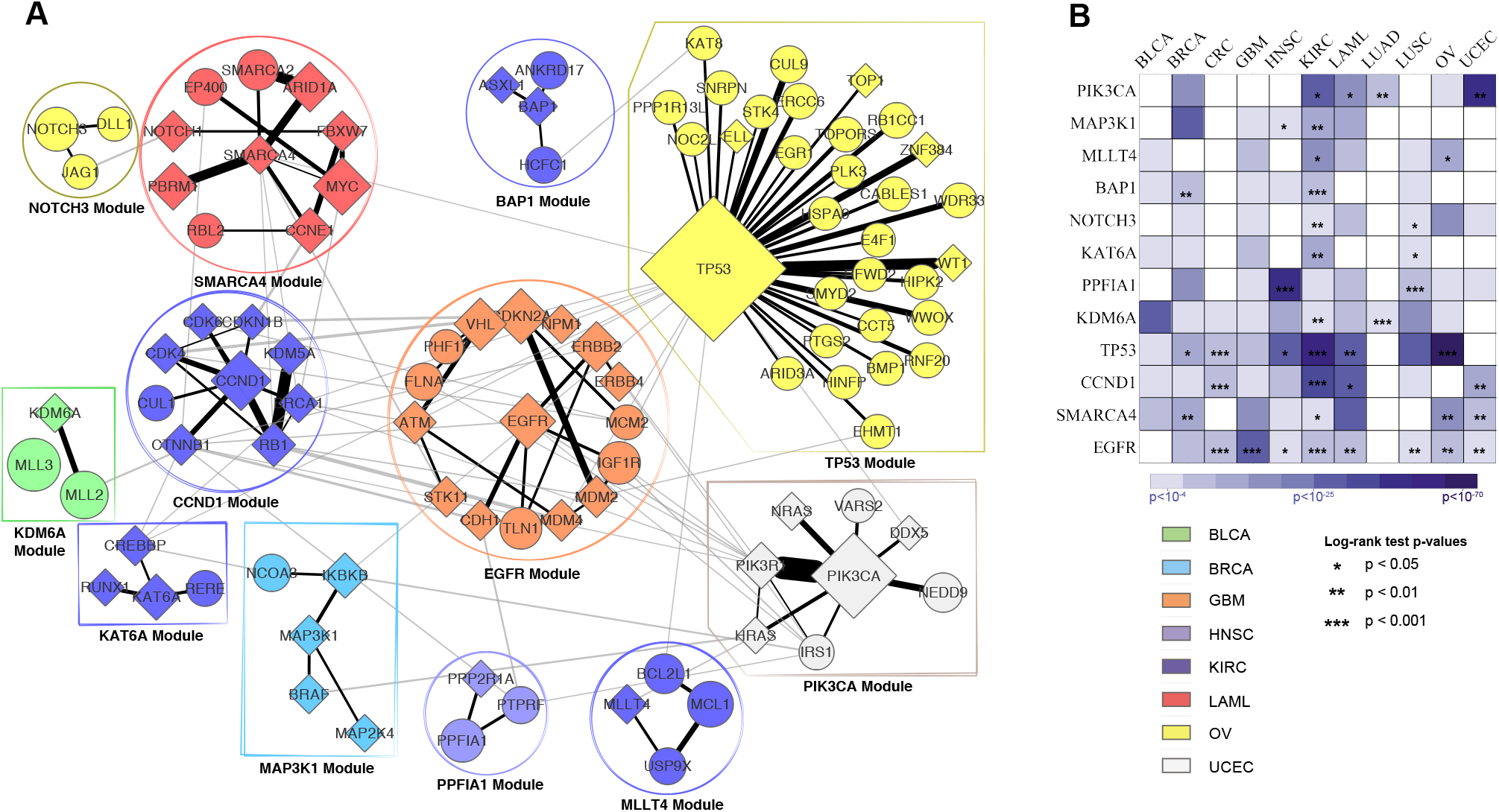
A) MEXCOwalk output modules when *total_genes* = 100. Diamond shaped nodes correspond to CGC genes. Sizes of the nodes are proportional with mutation frequencies of corresponding genes. Edges within the module are colored black, whereas the edges between the modules are colored. Edge weights are reflected in the thicknesses of the line segments. Color of a module denotes the cancer type with the strongest enrichment for mutations in genes of that module. The legend for the color codes are shown on the right. Each module is named with the largest degree gene in the module. B) Results of cancer type specificity and survival analyses. Rows correspond to modules and columns correspond to cancer types. Colors of the matrix entries indicate the significance of enrichment for cancer types in terms of Fisher’s exact test p-values. Stars indicate the significance of log-rank test p-values in survival analyses.

Many of these modules are part of well known cancer-related pathways such as those centered at EGFR, TP53, PIK3CA, and CCND1. Analyzing the interactions between the modules, EGFR module can be seen as an important hub module between some important modules such as the TP53 module, CCND1 module, and the PIK3CA module; without the EGFR module these three modules would almost be isolated in the induced subgraph. The EGFR module contains several known cancer genes many of which are related to cell cycle control: VHL, CDKN2A, NPM1, ERBB2, ERBB4, MDM2, MDM4, STK11, CDH1, ATM. Seven cancer types are enriched for mutations in this module with GBM being the most significant enrichment; Fisher’s exact test p-value is = 3.5*e* − 21. Indeed, EGFR gene is mutated in more than half of all GBM patients and anti-EGFR agents are already used for GBM treatment (Taylor *et al*., 2012). However, resistance to these agents is a major problem suggesting that treatment strategies might benefit from targeting multiple genes in this module. This module also contains TLN1, which is not one of the known cancer genes listed in CGC. However it is mutated in 104 patients across 10 cancer types and it has previously been associated with tumorigenecity and chemosensitivity (Singel *et al*., 2013; Fang *et al*., 2016). We investigate whether the genes in this module are predictive of patient survival profiles by calculating a risk score for each patient as in Beer *et al*. (2002) and Shrestha *et al*. (2017). When we divide the GBM patients into two as training and test sets, the low-risk and high-risk thresholds that we identify from the training set are successful in stratifying the patients into low-risk and high-risk groups in the test set with the log-rank test p-value= 0.0004; see Fig. 3-B. Our TP53 module includes 30 interactors out of 213 available in the HINT+HI2012 PPI network. TP53 shares the highest edge weight with WT1, which is a transcription factor that has roles in cellular development and cell survival. Another gene which has a large edge weight is CUL9. Its mutation frequency is only 0.015, which would possibly make it easy to miss through single-gene tests. The PIK3CA module identifies several genes in the PI3K pathway whose deregulation is critical in cancer development and progression (Karakas *et al*., 2006). The module provides a chance to observe the importance of incorporating mutual exclusivity in MEXCOwalk. Among all the interactions presented in the induced subgraph of 100 genes in all 12 modules, the one between PIK3CA and PIK3R has the largest weight. These genes are mutated in 602 and 155 patients respectively, although the overlap between the two patient sets is only 18 indicating the high mutual exclusivity between the pair of genes. The CCND1 module is yet another fairly known cancer driver module (Kim and Diehl, 2010; Malumbres and Barbacid, 2009). Other than EGFR, it is the module that contains the most reference genes; all 9 genes in the module except CUL1, are in the CGC database. It is shown that the mutations, amplification, and expression changes of these genes, which alter cell cycle progression, are frequently observed in a variety of tumors (Malumbres and Barbacid, 2009; Kim and Diehl, 2010). Indeed, we find significant association of this module with patients ‘ survival outcome in CRC, KIRC, LAML, and UCEC types; see Fig. 3-B.

A comparison of the output modules of Hotnet2 and MEMCover_v1 in the same setting of *total_genes* = 100 leads to interesting observations; see Supplementary Document for the plots. 47 genes are common between MEXCOwalk and MEMCover_v1, whereas only 32 genes are common between MEXCOwalk and Hotnet2. MEMCover_v1 identifies 76 modules in total. Out of these, 54 contain only a single gene and 20 contain two genes. We observe a similar result when we analyze MEMCover’s published results on HumanNet when *total_genes* is 100. Out of the 62 output modules, 31 are of size one and 27 are of size two indicating that this is not a bias we introduce by running MEMCover with a different dataset. With such a difference in module sizes, it is difficult to compare MEXCOwalk modules with those of MEMCover. Comparing modules of MEXCOwalk with those of Hotnet2, we observe several interactions between MEXCOwalk modules, whereas for Hotnet2, among all 100 genes, the only such interaction is between ATM and STK11. In total there are 48 genes of MEXCOwalk in the reference set, whereas Hotnet2 provides 28 such genes. Every MEXCOwalk module except the NOTCH3 module, contains a known driver. In contrast, 8 out of the 19 modules identified by Hotnet2 lack a known driver. Hotnet2 is unable to identify any of the genes in our CNND1 module which contains several cell cycle regulators which also include eight known cancer drivers. Similarly, the majority of the genes in our SMARCA4, MAP3K1, and EGFR modules containing several known drivers are not present among Hotnet2 modules.

## 4 Conclusion

In this study, we introduce a novel method, MEXCOwalk, that incorporates network connectivity, mutual exclusivity, and coverage information to identify cancer driver modules. MEXCOwalk outperforms several state-of-the-art computational methods on TCGA pan-cancer data in terms of recovering known cancer genes including the rarely mutated ones, providing modules that are capable of classifying normal and tumor samples, and that are enriched for mutations in specific cancer types. In the future, additional types of genetic and epigenetic aberrations can be incorporated as they become available. Finally, adaptations of MEXCOwalk to include network density-related scores in edge weights constitute planned extensions of this work.

## Acknowledgements

Authors are listed in alphabetical order with respect to lastnames.We thank Aissa Houdjedj for his help with the preparation of the figures.

