## Supplementary Material for "MEXCOwalk: Mutual Exclusion and Coverage Based Random Walk to Identify Cancer Modules"

#### 1 Proof of Theorem 1

**Theorem 1.** *Cancer driver module identification problem is NP-hard.*

*Proof.* The transformation is from *Set Packing* which is NP-complete [3]. In the Set Packing problem, given a collection $C$ of finite sets and a positive integer $K \leq |C|$, the problem is to find out whether $C$ contains at least $K$ mutually disjoint sets. The problem is NP-hard even when the size of each set is at most 3, which can easily be extended to the setting where the size of each set is exactly 3. Given an input to the Set Packing problem within this setting in the form of $K$ and $C$ such that for each $S \in C$, $|S| = 3$, we generate $G$ as a complete graph on $|C|$ vertices, corresponding to the such that each finite set in $C$ corresponds to a set of samples $S_i$ for which gene $g_i$ is mutated. We set both $total\_genes$ and $min\_module\_size$ to $K$. The answer to the Set Packing problem is Yes, if and only if the maximized score of the cancer module identification problem is exactly $\frac{3 \times K}{|\bigcup_{\forall g_i \in V} S_i|}$. □

#### 2 Effects of Mutual Exclusivity Threshold $\theta$

We plot the distribution of $MEX_n$ scores across all the edges in Figure S1. Almost all edges have $MEX_n$ scores larger than 0.5. Therefore, we experiment with $\theta$ values greater than or equal to 0.5: $\{0.5, 0.6, 0.7, 0.8, 0.9\}$. Figure S2 show the results of running MEXCOwalk with different $\theta$ values. In parts A and B, we observe that changing $\theta$ has a minimal effect in recovering known cancer genes with $\theta = 0.7$ giving the largest area under the ROC curve. Figure S2 -C reveals a strong correlation between $\theta$ scores and Mutual Exclusivity (MS) scores. Since large values of $\theta$ clamp a larger set of edge weights to 0, the resulting modules are only those with really large mutual exclusivity scores. We observe the largest coverage scores when $\theta$ is set to 0.7 (Figure S2-D), whereas 0.9 results in significantly lower coverage scores across all $total\_genes$ values. This is likely due to mutual exclusivity dominating over coverage in edge weights. Finally, we observe that Driver Module Set Scores are mosty in parallel with Coverage Scores; see Figure S2-B.

#### 3 Effects of *min_module_size*

We experiment with a range of values for *min_module_size*; see Figure S3. We see minimal differences in overlaps with CGC database where *min_module_size* = 3 results in the largest area under ROC curve. As expected, there is an inverse correlation between *min_module_size* and mutual exclusion scores (Figure S3-C). On the other hand, we observe a positive correlation between *min_module_size* and coverage scores. Again, we observe that Driver Module Set Scores are in parallel with Coverage Scores (Figure S3-B).

#### 4 Static Evaluations

Figure S4-A plots the Receiver Operating Characteristic (ROC) curves of the set of genes in the union of modules each algorithm provides with respect to the CGC genes [2]. In Figure S4-B and Figure S4-C, ROC curves are plotted with a subset of CGC genes with mutation frequency $\leq 1\%$ and $\leq 2\%$ in the pan-cancer cohort, respectively. The same analysis is repeated with the set of druggable genes downloaded from DGIdb 3.0 [1]. Note that only cancer related sources are included in DGIdb 3.0. FigureS5-A shows the ROC curve with all druggable genes; whereas in Figure S5-B and Figure S5-C only those genes with mutation frequency $\leq 1\%$ or $\leq 2\%$ are included, respectively.

#### 5 Output Module Sizes

We investigate the size distribution of the output modules for different methods under consideration. In Figure S8-A we plot the average module sizes for different *total_genes* values. We observe that the average module size remains stable for increasing values of *total_genes* for all the methods except Hotnet2. Additionally, we identify the minimum and maximum module size for each *total_genes* value and for each method. We then plot the percentage of the genes that belong to the modules of minimum or maximum size; see Figure S8-B and Figure S8-C. Strikingly, for the majority of *MEMCover_v1* outputs, more than 70% of genes appear in modules with minimum size which is 1. In the other extreme, Hotnet2 provides a large module of maximum size that contains more than 60% of all the genes, for the majority of *total_genes* settings.

#### 6 Cancer Type Specificity

Figure S10 shows the distribution of classification accuracy values across the output modules for increasing values of *total_genes* for all the methods.

### 7 Classification Accuracy

Figure S9 shows the distribution of best p-values obtained for each module for increasing values of *total_genes* for all the methods.

### 8 Hotnet2 and MEMCover output modules

Hotnet2 and MEMcover output modules are shown in Figure S11 when *total_genes* is set to 100. The module colors represent the mostly enriched cancer type. Note that in Figure S11-B, every single gene not surrounded by a rectangle is a module by itself.

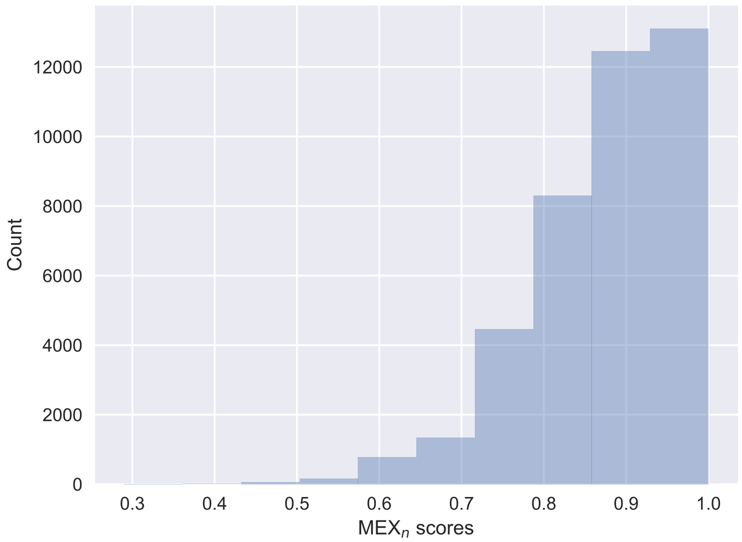


Figure S1: The distribution of $MEX_n$ scores of all edges.

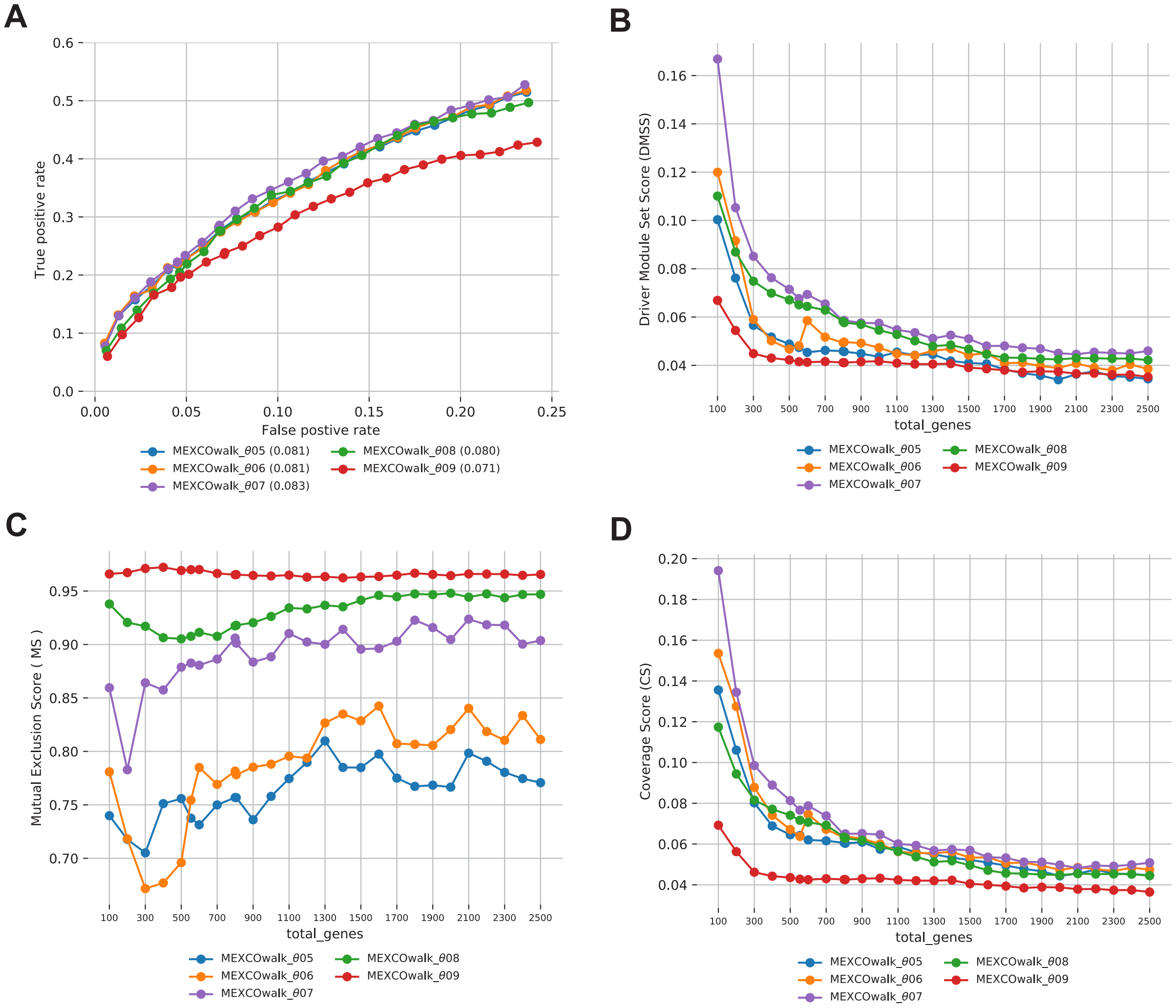


Figure S2: Comparison of MEXCOwalk models with different mutual exclusion score thresholds ($\theta$): 0.5, 0.6, 0.7, 0.8 and 0.9 A) ROC plots and AUROC values written in parentheses. B) Driver Modules Set Score (DMSS). C) Mutual Exclusion Score (MS). D) Coverage Score (CS).

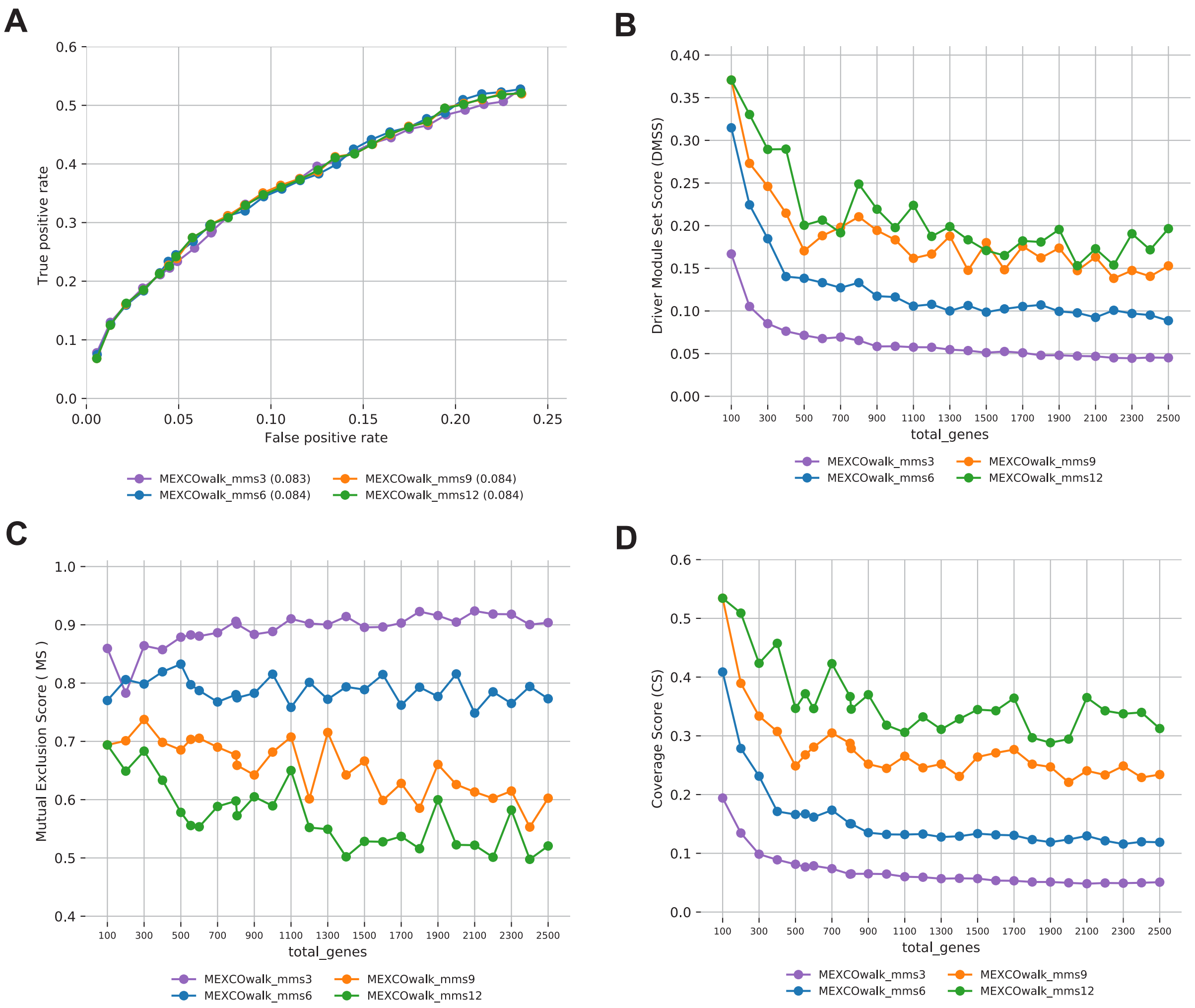


Figure S3: Comparison of MEXCOwalk models with different $min\_module\_size$: 3, 6, 9, 12 A) ROC plots and AUROC values written in parentheses. B) Driver Modules Set Score (DMSS). C) Mutual Exclusion Score (MS). D) Coverage Score (CS).

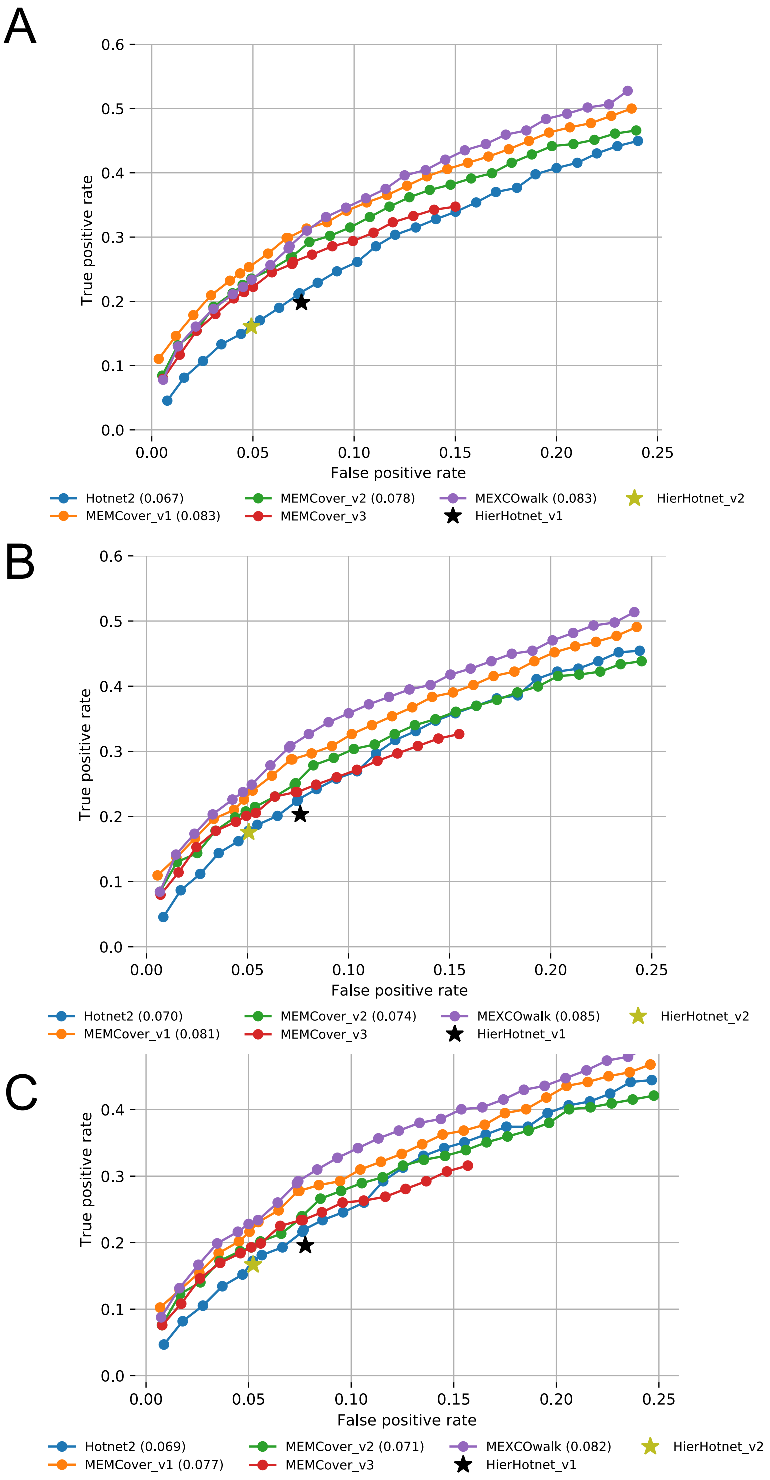


Figure S4: A) The fraction of recovered CGC genes for each *total_genes* value is shown with a ROC plot. B) Same as A, but only those CGC genes with $\leq$ 1% mutation frequency in the pan-cancer cohort are used. C) Same as A, but only those CGC genes with $\leq$ 2% mutation frequency in the pan-cancer cohort are used.

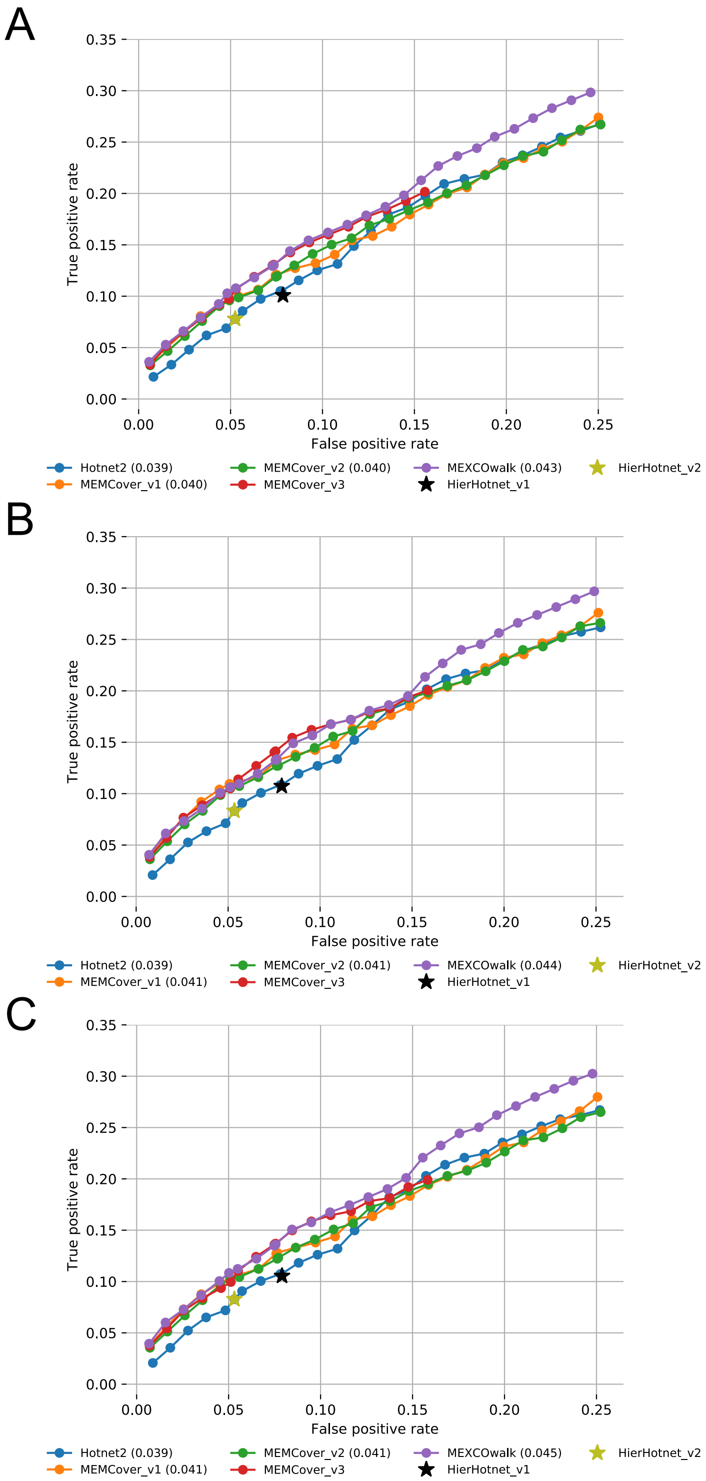


Figure S5: A) The fraction of recovered druggable genes for each *total_genes* value is shown with a ROC plot. B) Same as A, but only those druggable genes with $\leq 1\%$ mutation frequency in the pan-cancer cohort are used. C) Same as A, but only those druggable genes with $\leq 2\%$ mutation frequency in the pan-cancer cohort are used.

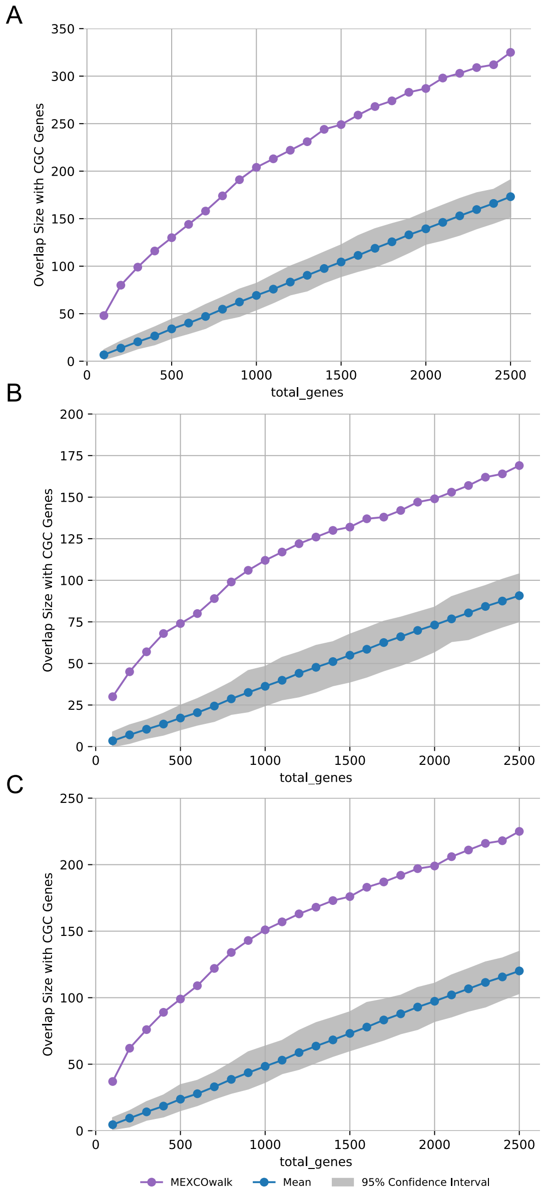


Figure S6: A) Overlap size CGC genes for each *total_genes* value is shown with a ROC plot for MEXCOWalk on original and randomized mutation data. B) Same as A, but only those CGC genes with $\leq$ 1% mutation frequency in the pan-cancer cohort are used. C) Same as A, but only those CGC genes with $\leq$ 2% mutation frequency in the pan-cancer cohort are used.

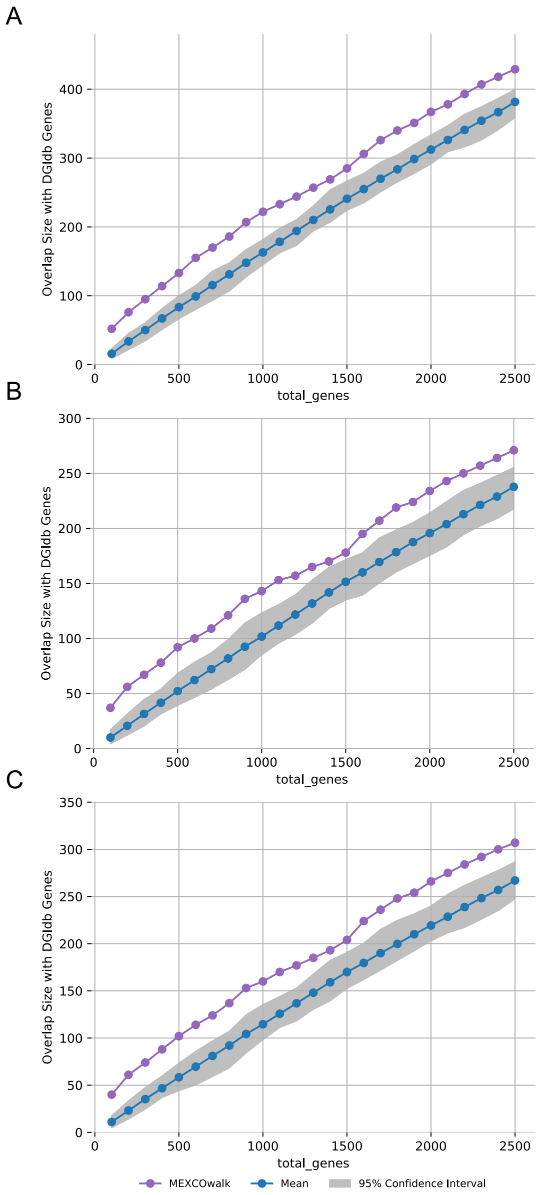


Figure S7: A) Overlap size with recovered druggable genes for each *total_genes* value is shown with a ROC plot for MEXCOWalk on original and randomized mutation data. B) Same as A, but only those druggable genes with $\leq 1\%$ mutation frequency in the pan-cancer cohort are used. C) Same as A, but only those CGC genes with $\leq 2\%$ mutation frequency in the pan-cancer cohort are used.

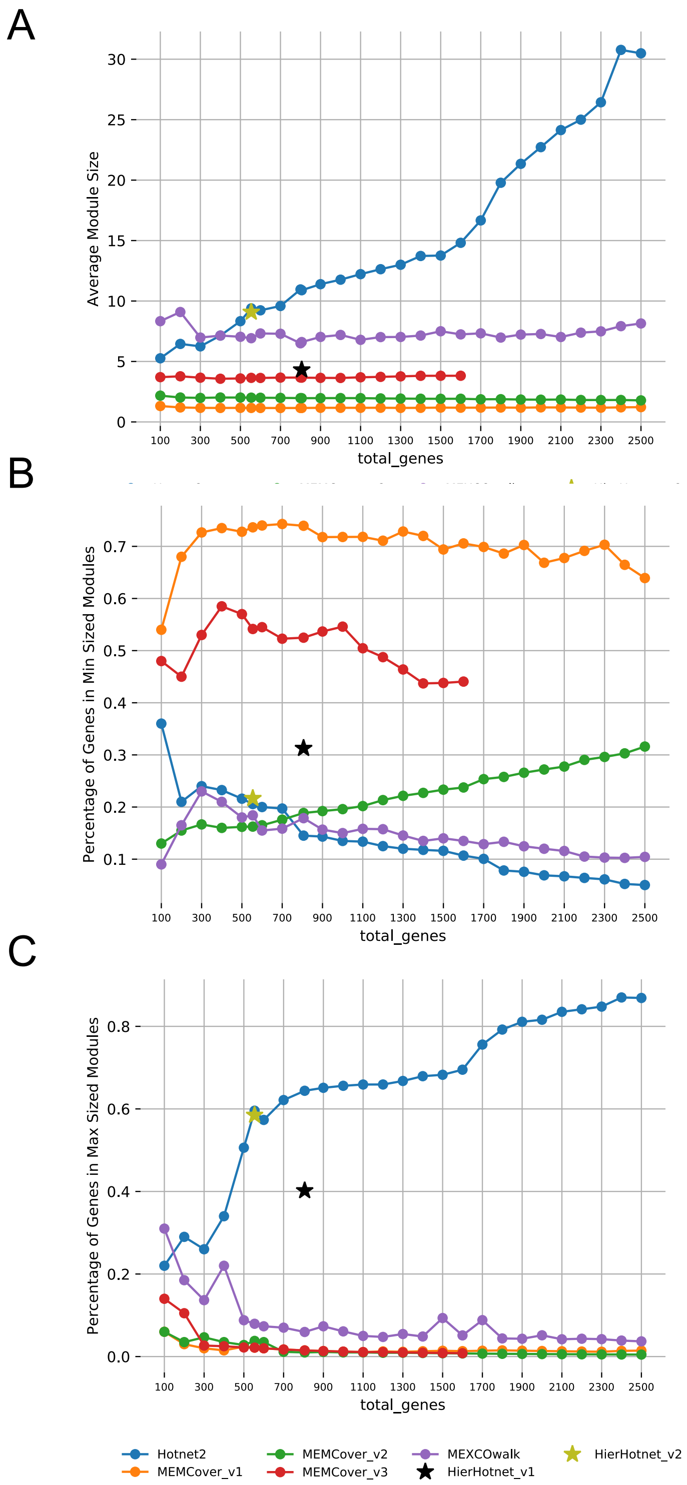


Figure S8: A) Average modules sizes in the outputs of MEXCOwalk, MEMCover, Hotnet2, and Hierarchical Hotnet for increasing values of *total_genes*. B) The percentage of genes across the modules with largest size for MEXCOwalk, MEMCover, Hotnet2, and Hierarchical Hotnet outputs for increasing values of *total_genes*. C) The percentage of genes across the modules with smallest size for MEXCOwalk, MEMCover, Hotnet2, and Hierarchical Hotnet outputs for increasing values of *total_genes*.

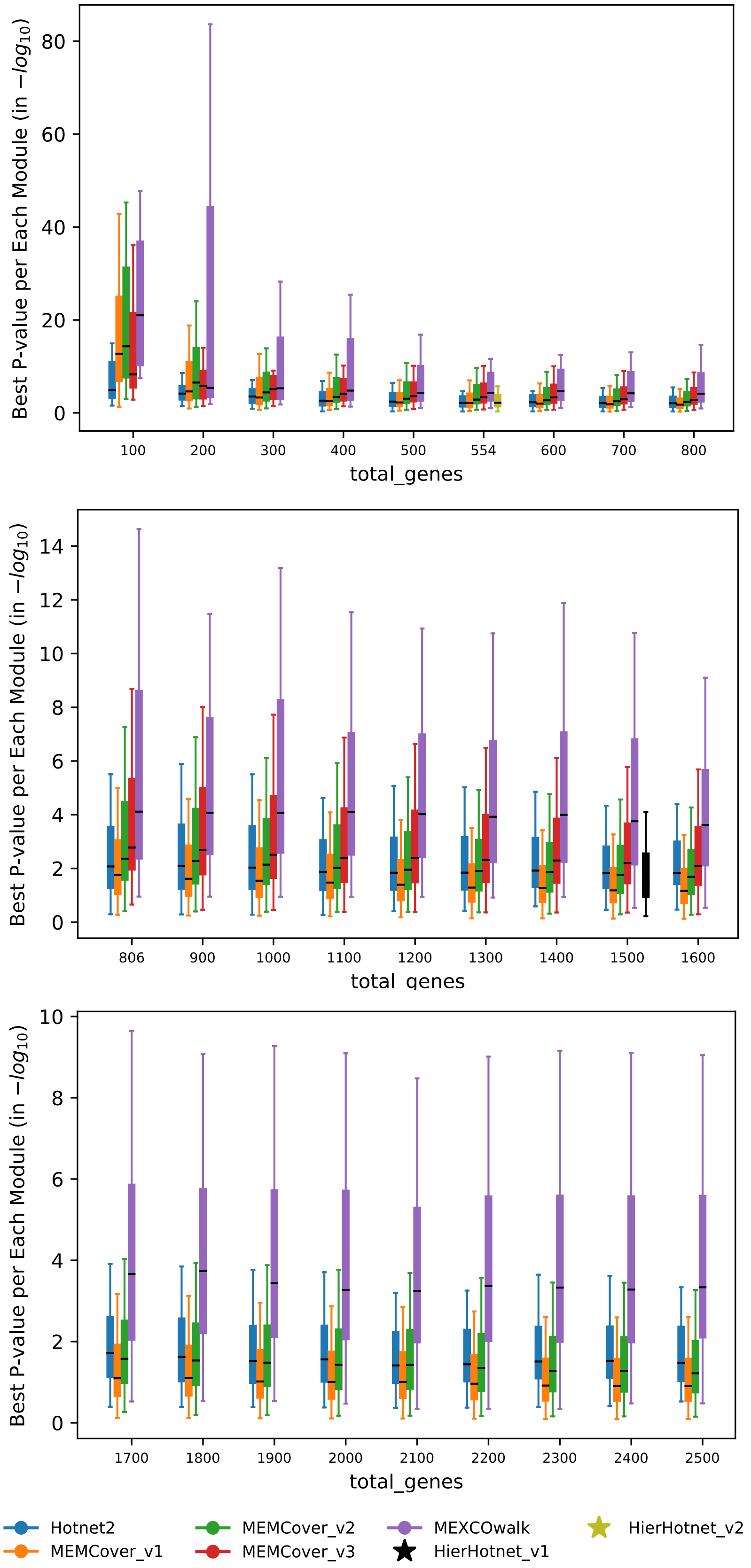


Figure S9: Box plot figure of best p-value for each module obtained for increasing values of *total_genes*. Whiskers represent the interquartile range.

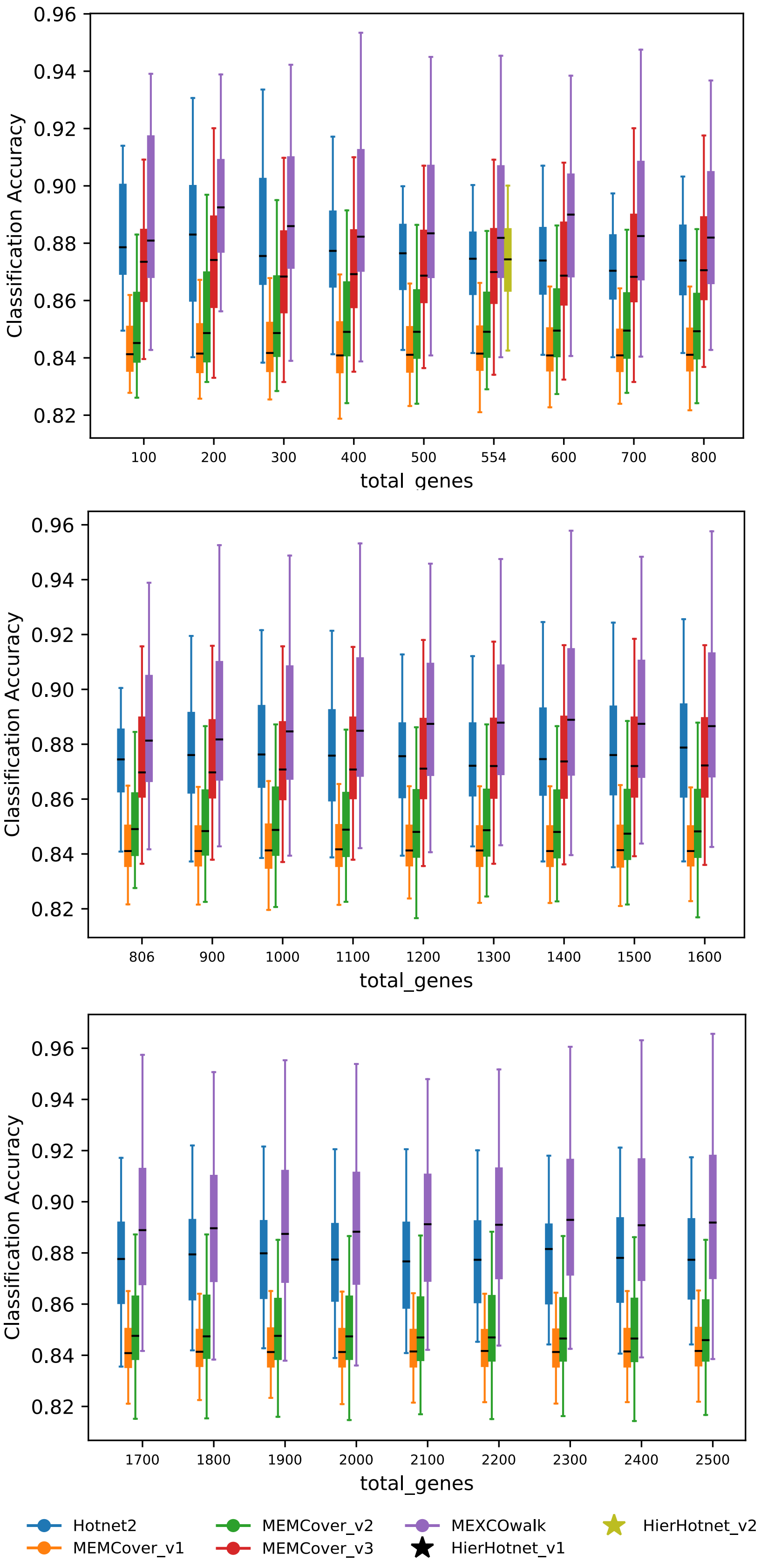


Figure S10: Box plot figure of classification accuracy for each module obtained for increasing values of *total_genes*. Whiskers represent the interquartile range.

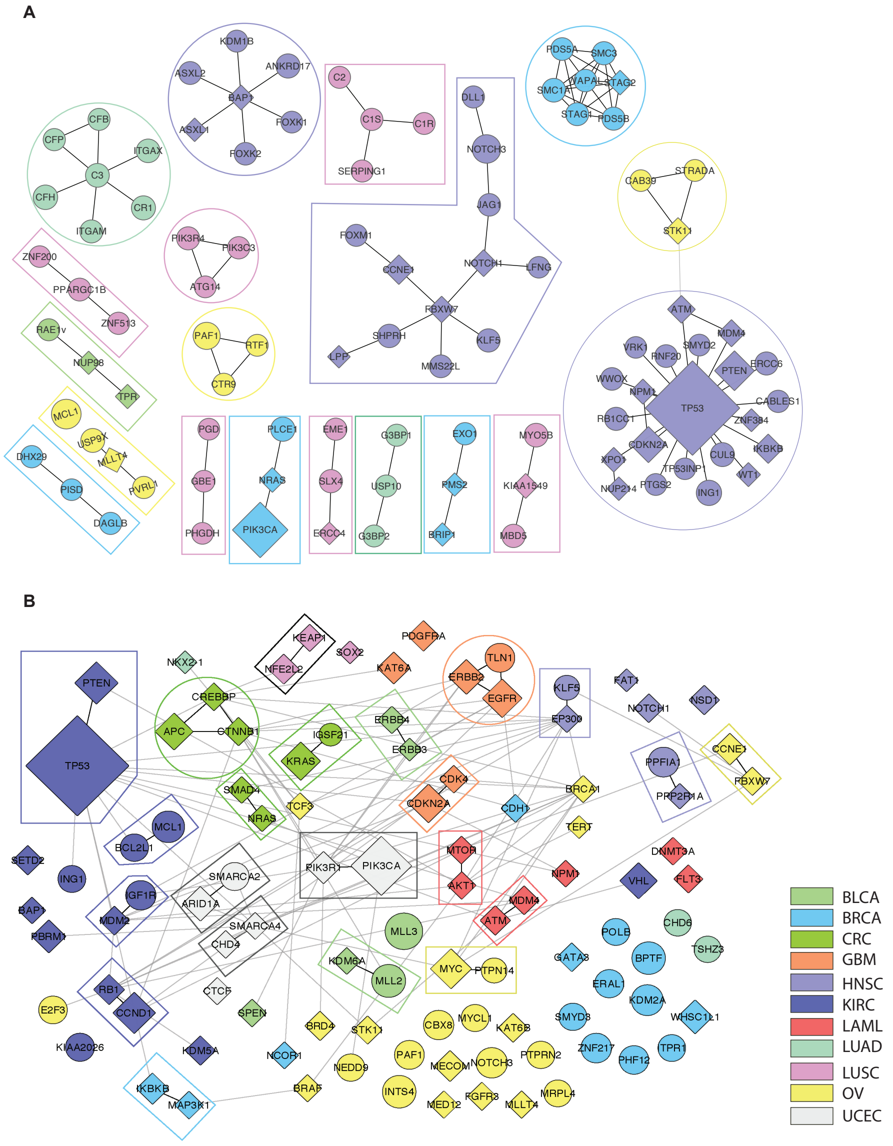


Figure S11: A) Hotnet2 output modules when $total\_genes = 100$. B) MEMCover_v1 output modules when $total\_genes = 100$ Diamond shaped nodes correspond to CGC genes. Sizes of the nodes are proportional with mutation frequencies of corresponding genes. Color of a module denotes the cancer type with the strongest enrichment for mutations in genes of that module. The legend for the color codes are shown on the bottom right.
